# Insights into post-translational regulation of skeletal muscle contractile function by the acetyltransferases, p300 and CBP

**DOI:** 10.1101/2024.02.27.582179

**Authors:** Gretchen A. Meyer, Jeremie L.A. Ferey, James A. Sanford, Liam S. Fitzgerald, Akiva E. Greenberg, Kristoffer Svensson, Michael J. Greenberg, Simon Schenk

**Affiliations:** Program in Physical Therapy; Departments of Neurology, Orthopaedic Surgery and Biomedical Engineering and; Department of Biochemistry and Molecular Biophysics, Washington University School of Medicine, St. Louis, MO, USA; Biological Sciences Division, Pacific Northwest National Laboratory, Richland, WA, USA; Department of Orthopaedic Surgery; Biomedical Sciences Graduate Program and; Medical Scientist Training Program, School of Medicine, University of California San Diego, La Jolla, CA, USA

**Keywords:** physiology, mitochondria, actin, myosin, force

## Abstract

Mice with skeletal muscle-specific inducible double knockout of the lysine acetyltransferases, p300 (E1A binding protein p300) and CBP (cAMP-response element-binding protein binding protein), referred to as i-mPCKO, demonstrate a dramatic loss of contractile function in skeletal muscle and ultimately die within 7 days. Given that many proteins involved in ATP generation and cross-bridge cycling are acetylated, we investigated whether these processes are dysregulated in skeletal muscle from i-mPCKO mice and thus could underlie the rapid loss of muscle contractile function. Just 4-5 days after inducing knockout of p300 and CBP in skeletal muscle from adult i-mPCKO mice, there was ∼90% reduction in *ex vivo* contractile function in the extensor digitorum longus (EDL) and a ∼65% reduction in *in vivo* ankle dorsiflexion torque, as compared to wildtype (WT; i.e. Cre negative) littermates. Despite the profound loss of contractile force in i-mPCKO mice, there were no genotype-driven differences in fatigability during repeated contractions, nor were there genotype differences in mitochondrial specific pathway enrichment of the proteome, intermyofibrillar mitochondrial volume or mitochondrial respiratory function. As it relates to cross-bridge cycling, remarkably, the overt loss of contractile function in i-mPCKO muscle was reversed in permeabilized fibers supplied with exogenous Ca^2+^ and ATP, with active tension being similar between i-mPCKO and WT mice, regardless of Ca^2+^ concentration. Actin-myosin motility was also similar in skeletal muscle from i-mPCKO and WT mice. In conclusion, neither mitochondrial abundance/function, nor actomyosin cross-bridge cycling, are the underlying driver of contractile dysfunction in i-mPCKO mice.

**New & Noteworthy:** The mechanism underlying dramatic loss of muscle contractile function with inducible deletion of both p300 and CBP in skeletal muscle remains unknown. Here we find that impairments in mitochondrial function or cross-bridge cycling are not the underlying mechanism of action. Future work will investigate other aspects of excitation-contraction coupling, such as Ca^2+^ handling and membrane excitability, as contractile function could be rescued by permeabilizing skeletal muscle, which provides exogenous Ca^2+^ and bypasses membrane depolarization.

## Introduction

Reversible acetylation of lysine residues is a common and conserved post-translational modification that contributes to a diverse array of cellular processes, including energy metabolism, transcription and cell cycle progression (1). The full extent of physiological regulation via lysine acetylation is only beginning to be appreciated with the refinement of mass spectrometry and application of unbiased proteomics. In skeletal muscle, acetyl-lysine-specific proteome analysis demonstrates that the prevalence of lysine acetylation is comparable to other major posttranslational modifications such as phosphorylation and ubiquitylation (2). Indeed, proteins within essentially all aspects of skeletal muscle homeostasis and function are acetylated, including proteins contributing to substrate metabolism, mitochondrial electron transport, transcription, excitation-contraction (E-C) coupling and contractile function (2, 3). Despite this prevalence, the contributions of lysine acetylation to skeletal muscle physiology and function are largely unknown.

Generally speaking, acetylation of a given lysine residue is a balance between the actions of acetyltransferases, which add an acetyl group from acetyl coenzyme A (CoA) and deacetylases, which remove the acetyl group. In cells from humans and mice, there are ∼22 known acetyltransferases spread across three main families, general control of amino acid synthesis 5 (GCN5), E1A binding protein p300 (p300)/cAMP response element-binding protein-binding protein (CBP) and MYST (1). To date, the contribution of acetyltransferases to cellular biology has largely focused on their presence in the nucleus and their role in the regulation of transcription, such as through their regulation of histone acetylation. Interestingly, however, unlike most acetyltransferases, p300/CBP also have a strong presence in the cytoplasm, as well as the nucleus (4-6), including in skeletal muscle (7). Indeed, in human platelets, which have no nucleus, p300 directly regulates lysine acetylation of many cytosolic proteins and consequently, many aspects of platelet biology and function (6). Thus, the contribution of p300/CBP to cellular biology clearly extends to the cytosol and beyond their established role in the nucleus.

Recently, we established a critical role for p300 and CBP in the maintenance of skeletal muscle contractile function (7). Mice with inducible, skeletal muscle-specific double knockout of p300 and CBP (i-mPCKO) demonstrate, in just 3-5 days, overt loss of transcriptional homeostasis, a dramatic loss of contractile function and ultimately mortality (7). While this work demonstrated that p300 and CBP are necessary for normal muscle contractile function, it did not establish potential mechanisms of regulation. For example, in keeping with the well-documented nuclear role of p300/CBP, we saw profound changes in transcriptional control and gene expression in i-mPCKO mice, especially of genes annotating to various aspects of E-C coupling and contractile function (7). However, we found minimal changes in protein abundance, which would be expected to underlie functional changes (7), suggesting that a non-transcriptional mechanism likely mediates the effects of p300/CBP on skeletal muscle contractile function. Considering p300/CBP are present and functionally important in the cytoplasm, we hypothesized that they may regulate contraction via non-nuclear mechanisms, including through actions in the cytoplasm. Supporting this, recent acetylomics analysis identified two primary categories of acetylated proteins in skeletal muscle cytoplasm: metabolic proteins involved in the generation of ATP and sarcomeric proteins supporting cross-bridge cycling (2). A functional role for p300/CBP in metabolic regulation is supported by our work demonstrating a role for p300/CBP in the regulation of insulin-stimulated glucose uptake (8), whilst a functional role for acetylation in cross-bridge cycling is supported by work in rabbit and drosophila muscle demonstrating modulation of contractile force by acetylation of troponin and actin, respectively (9-11). Considering evidence of a potential non-nuclear role for p300/CBP in the regulation of muscle contractile function, the goal of this work was to evaluate the contribution of mitochondrial function and cross-bridge cycling to the contractile phenotype in i-mPCKO mice, and by extension, to home in on potential mechanisms by which p300/CBP regulate skeletal muscle contractile function. We hypothesized that p300/CBP acyltransferase activity is required for skeletal muscle contractile function, and that this regulatory action is due, in part, to impairments in myosin and actin cross-bridge cycling and mitochondrial respiratory capacity.

## Materials and Methods

### Study approval

All animal work described was performed in accordance with the National Institutes of Health’s Guide for the Use and Care of Laboratory Animals and was approved by the Animal Studies Committee of the Washington University School of Medicine and the Animal Care Program and Institutional Animal Care and Use Committee at the University of California, San Diego.

### i-mPCKO mouse model

The establishment of the i-mPCKO mouse line has been described in detail elsewhere (7, 8). Briefly, i-mPCKO mice were generated through judicious interbreeding of tamoxifen (TMX)-inducible, human α-skeletal actin (HSA)-Cre recombinase mice (12) with *Ep300* and *Crebbp* floxed mice (LoxP sites flanking exon 9 of both genes; (13, 14)). i-mPCKO mice (i.e. p300^flox/flox^ and CBP^flox/flox^ and Cre positive [Cre^+^; on one allele]) and wildtype (WT) mice (i.e. p300^flox/flox^ and CBP^flox/flox^ and Cre negative [Cre^–^])were bred together to generate study mice; this strategy was used as it generated both Cre^+^ (i.e. i-mPCKO) and Cre^–^ (i.e. WT) littermates.

### Experimental design

3-6 month old mice were housed in a temperature controlled (22°C) facility with a 12 hour light-dark cycle with *ad libitum* access to standard chow. Depending on the experiment, both male and female mice were studied, as noted in figure legends. All experiments and tissue collection were performed between 08:00 and 15:00, during the light phase. For studies in i-mPCKO mice and WT littermates, prior to experimentation, all mice were dosed daily with tamoxifen (TMX; 2 mg) by oral gavage or intraperitoneal (i.p.) injection for up to five consecutive days; day 0 (D0) indicates the first day of TMX dosing, while day 5 (D5) is the day after the last TMX treatment.

### Ex-vivo muscle contractility

*Ex-vivo* contractile function was assessed in the 5^th^ toe belly of the extensor digitorum longus (EDL) muscle of WT and i-mPCKO mice at D5 post TMX as previously described (7). Briefly, muscles were mounted to a dual-mode force-length transducer and static pin in a chamber filled with Ringer’s solution at 25°C. Prior to stimulation, muscle length was increased until sarcomere length reached optimal (2.1-2.4 μm) at which point fiber length was measured through a microscope. Then supramaximal conditions were set for each muscle by gradually increasing current until twitch force plateaued and then increasing this value by 50%. For contractile function measurements, muscles were stimulated at their supramaximal current through parallel platinum electrodes at 100 Hz (300ms with 0.5ms pulse width). In a subset of trials, fatiguability was also measured by eliciting tetanic contractions every 15 seconds for 10 minutes. Following testing, muscles were removed and weighed. Force was normalized to muscle physiological cross-sectional area, as described (15), for all comparisons.

### Serial measurement of in-vivo muscle contractility

In i-mPCKO and WT littermates, ankle dorsiflexion torque was measured via a custom physiology rig (Aurora Scientific; 1300A). Mice were deeply anesthetized with 2% inhaled isoflurane and hindlimbs shaved and sterilized prior to transfer to the pre-warmed rig platform. The hindlimb was then clamped at the knee and the foot was secured to a footplate attached to a dual-mode force-length transducer such that the ankle was positioned at 90° with the tibia parallel to the floor. Sterile 27-gauge needle electrodes were inserted through the skin over the tibialis anterior (TA) muscle near the peroneal nerve endplates, taking care to avoid muscle puncture. Following electrode insertion, twitch and tetanic (20V, 100Hz, 300ms with 0.3ms pulse width) contractions were recorded. Serial recordings were made on D0 (prior to TMX treatment), D2 and D4 post TMX. Following recordings on D4, fatiguability was also measured by eliciting tetanic contractions every 10 seconds for two minutes.

### Permeabilized fiber testing

EDL muscles were dissected from WT and i-mPCKO mice at D5 post TMX, trimmed to remove tendons and permeabilized in a solution of Brij 58 (0.5% w/v) with agitation for 30 minutes. Muscles were then stored in a glycerinated storage solution (50% w/v) at -20°C until testing; muscles were tested within 2 months of muscle collection. Active tension was measured in individual fibers over a range of Ca^2+^ concentrations as previously described in detail (16). Briefly, a 2-3 mm segment of permeabilized myofiber was teased from the muscle bulk in a pCa 9 (8mM ATP, 10mM creatine phosphate) relaxing solution and affixed in a chamber within the Aurora permeabilized fiber system (Aurora Scientific, 1400A) bathed in the same solution with one end tied to a lever arm and the other to a force transducer with 10-0 nylon suture. An inverted light microscope was used to set sarcomere length to 2.5 μm and to measure fiber diameter at three places along the length of the fiber. The fiber was then transferred to a weakly buffered pre-activating solution followed by a pCa 4.5 (8mM ATP, 10mM creatine phosphate) activating solution by chamber switching in the 1400A system. This was repeated 3 times. At the end of active testing, fibers were returned to relaxing solution to equilibrate for 5 minutes. Then, fiber activation was similarly measured in 8 solutions of varying Ca^2+^ concentration, from pCa 4.5 to pCa 6 in a randomized order. All forces were normalized to fiber cross-sectional area, calculated from the average fiber diameter and adjusted for the swelling effects of permeabilization (17). Force-pCa data were fit with a Hill equation to calculate EC_50_ (18).

### Mitochondrial Pathway Analysis

We mined publicly available quantitative proteomics data in muscle from WT mice (D5 post TMX) and i-mPCKO mice (D3 and D5 post TMX) from the PRoteomics IDEntifications Database (PRIDE project: PXD015694), which we previously generated (7). Pathways from the MitoCarta 3.0 database (19) were tested by fast gene set enrichment analysis (FGSEA) to identify mitochondrial pathways differentially enriched in wild-type and transgenic muscle. FGSEA was performed with *fgsea::fgseaMultiLevel* (v1.28.0) in R. Prior to analysis, MitoCarta pathways were filtered to those with at least 50% of their members detected in our dataset, resulting in 89 pathways for testing. Ranking metrics for each protein were generated using the statistics included in the PRIDE release, and were calculated as the signed-log_10_-transformed p-value, where the sign indicates the direction of the log_2_ fold change. For preliminary estimation of enrichment p-values and calculation of normalized enrichment scores (NES), a total of 10,000 permutations were used, and p-values were adjusted across results from all comparisons using the BH procedure. Heatmaps of the enrichment results display the pathways with the highest -log10-transformed adjusted p-value, and circles are scaled by p-value. Circles are colored by NES, and NES values between -1 and 1 are not shown. For plots of individual proteins comprising complex I (CI) and complex II (CII) of the electron transport chain, intensity values from the proteomics dataset were ratioed to the average intensity of the 3 WT mice. Proteins are labeled by UniProt symbol.

### Transmission Electron Microscopy

Transmission electron microscopy images were acquired from the EDL of i-mPCKO and WT mice at D5 post TMX, as previously described (7). In images from this work, the intermyofibrillar mitochondrial volume was assessed by point counting (20). The grid spacing was 0.3 *μ*m along both x- and y-axes. Mitochondria boundaries were recognized at ×5000 magnification and identified as intermyofibrillar as mitochondria that were separated from the sarcolemma by at least one myofibril. Intermyofibrillar mitochondrial volume was calculated by dividing the points assigned to mitochondria by the total number of points counted. Data are the average of 10 images per muscle.

### High-resolution respirometry

The EDL was excised from mice at D5 post TMX, and fibers were separated in BIOPS solution (Oroboros Instruments) on ice, permeabilized with saponin (0.05 mg/ml) for 20 min at 4° C, washed, blotted dry, and then weighed. One to two milligrams of tissue was placed in an Oxygraph 2K (Oroboros Instruments) chamber containing 2 ml of MirO5 (Oroboros Instruments) with 20 mM creatinine monohydrate and 10 μM blebbistatin. For the substrate-uncoupler-inhibitor titration (SUIT) protocol to measure carbohydrate-driven O_2_ flux, the following substrates were added sequentially, and measurements were performed at steady state (final concentrations indicated): leak: 0.5 mM malate, 5 mM pyruvate, 10 mM glutamate; complex I: 5 mM ADP; complex I+II: 10 mM succinate; and complex II: 0.5 μM rotenone. Samples were assayed in random order by a blinded investigator.

### Actin/myosin motility assay

Skeletal muscle myosin and actin were simultaneously prepared from limb skeletal muscles of 3 mice that were freshly excised, pooled and frozen in liquid nitrogen. Two separate preparations of n=3 pooled mice (WT and i-mPCKO mice at D5 post starting of TMX dosing) were run. Myosin was purified using the method of Kielley and Bradley (21). Full length myosin was stored in 0.5 M KCl, 10 mM EDTA pH 7.0, and 10 mM DTT at -80 °C. Skeletal actin was simultaneously purified from the same tissue using the method of Spudich and Watt (22) as previously described (23). *In* vitro motility was assessed in KMg25 buffer (25 mM KCl, 1 mM DTT, 2 mM EGTA, 4 mM MgCl_2_, and 60 mM MOPS pH 7.0) at 23°C as previously described (23, 24), unless noted otherwise. Briefly, myosin deadheads were removed by ultracentrifugation in the presence of 1 μM phalloidin-stabilized actin and 2 mM Mg.ATP before myosin was diluted to 140 μg/mL. Actin was polymerized on ice and filaments were stabilized with a molar equivalent of rhodamine labeled phalloidin. ATP concentrations were determined spectroscopically before each experiment. Flow chambers were constructed between two glass coverslips and the following solutions were sequentially added to the flow chamber: 100 μg/mL myosin in high salt buffer; 1 mg/mL bovine serum albumin in KMg25 (2 volumes); 1 μM phalloidin stabilized actin in KMg25; KMg25 + 5 mM Mg.ATP (2 volumes); KMg25 (4 volumes); 30 nM rhodamine phalloidin stabilized actin in KMg25 (1 volume); 1 volume activation buffer (KMg25 plus 192 U/mL glucose oxidase, 48 μg/mL catalase, 1 mg/mL glucose, 5 mM Mg.ATP, and 0.5% methyl cellulose). Motilily was imaged using a Retiga camera on an inverted microscope. Data were collected at 1 second per frame for 1 minute. 48-50 filaments for each condition were manually tracked using ImageJ with the MTrackJ Plugin. Filament speeds were normalized to the mean speed of WT for each run.

### Statistical analyses and reproducibility

For statistical analysis, data from male and female mice were combined as no differences in muscle contractile phenotype were uncovered between sexes (7). Between group comparisons were made by Student’s t-test or 2-way ANOVA as indicated, with Šídák’s multiple comparisons test. Actual numbers per group are indicated in each figure for each analysis. All results are presented as mean ± standard deviation. All statistical analyses were performed with GraphPad Prism. All chemicals were purchased from Fisher Scientific.

## Results

### Severely impaired ex vivo and in vivo muscle contractile function, but unaffected fatiguability, in i-mPCKO mice

Consistent with our previous work (7), i-mPCKO mice demonstrated dramatically reduced (– 90%) *ex vivo* contractile tension in the EDL at D5 post TMX, as compared to WT littermates (Fig 1A). Interestingly, however, EDL fatigability was similar between genotypes, (Fig 1B); from this data, the initial linear rate of fatigue (0-200 sec; Fig 1C) and final equilibrium relative tension values (Fig 1B) were also not different between groups. To investigate whether this contractile dysfunction in i-mPCKO mice existed *in vivo*, we examined ankle dorsiflexor torque and fatiguability through stimulation of the peroneal nerve in the same mice over a 5-day period. Peak dorsiflexion torque before initiating TMX dosing (day 0 [D0]) was not different between i-mPCKO and WT, nor was it different at D2 (Fig 1D). However, by D4 *in vivo* dorsiflexion torque was decreased 67±17% from D0 (Fig 1D). Despite these differences in *in vivo* contractile function, the linear rate of fatigue of dorsiflexion torque during a fatigue test was not significantly different between genotypes (Fig 1E and 1F).

**Figure 1.**
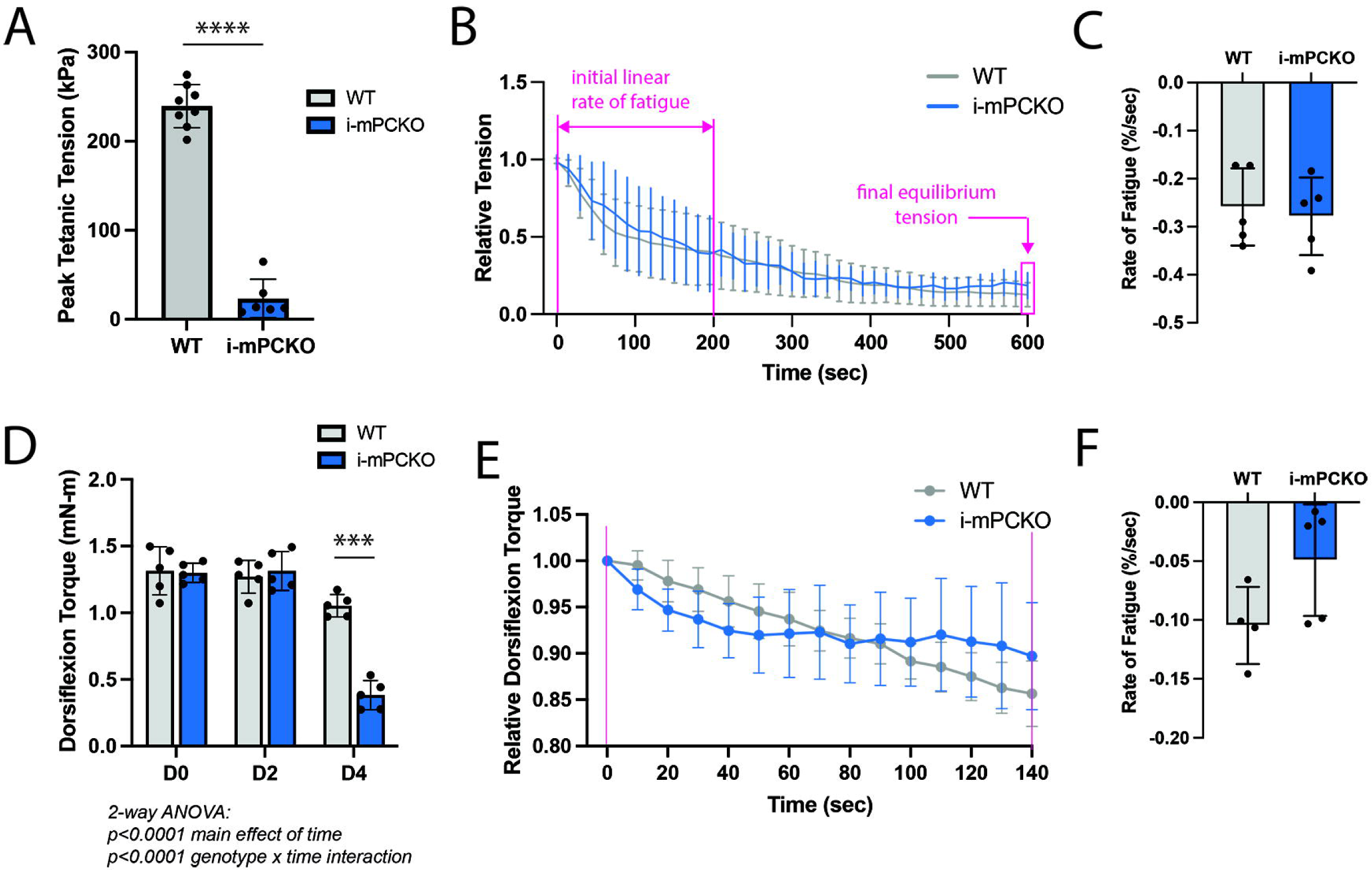
Both *ex vivo* and *in vivo* contractile function are dramatically impaired in i-mPCKO mice, with no effect on fatigability. ***(A)*** Peak tetanic tension measured *ex vivo* in isolated EDL muscles (D5 post TMX; data are for male and female, which were collapsed together, for each genotype). ***(B)*** Rate of relative tension decline during a fatiguing bout of contractions (D5 post TMX; only male mice only). The range for assessment of initial rate of fatigue and final equilibrium tension are marked in pink. ***(C)*** Quantification of the initial rate of fatigue of the data from *(B),* as the slope of the fatigue curves between 0-200 seconds. ***(D)*** Peak ankle dorsiflexion torque in the same mice, measured *in vivo* via stimulation of the peroneal nerve at D0, D2 and D4 after starting TMX dosing; data are for male and female, which were collapsed together, for each genotype. (D) Rate of decline in relative dorsiflexion torque during a fatiguing bout of repeated contractions on D4 post TMX; data are for male and female, which were collapsed together, for each genotype. The range for assessment of initial rate of fatigue is marked in pink. (E) Quantification of the initial rate of fatigue from data in *(D)*, calculaed as the slope of the fatigue curves between 0-140 seconds. Statistics: (A, C and F) Student’s unpaired t-test **** p<0.0001. D, 2-way ANOVA with repeated measures across time, Šídák’s multiple comparisons test *** p<0.001.

### Mitochondrial enrichment, abundance and function in skeletal muscle are not different between i-mPCKO and WT mice

In our publicly available proteomics dataset (7), no mitochondrial specific pathways from the MitoCarta 3.0 database were significantly differentially enriched between WT and i-mPCKO (Fig 2A; adjusted P>0.05). Also, of the 53 proteins comprising CI and CII that we were detected in the proteomic dataset, the abundance of only 1 (NDUFA11 protein that comprisein CIand CII was different between genotypes; the relative protein abundance for 10 of these proteins is plotted in Fig 2B, including NDUFA11. Intermyofibrillar mitochondria in the EDL appeared morphologically similar between WT and i-mPCKO (Fig 2C). Quantification of intermyofibrillar mitochondrial volume also found no differences between WT and i-mPCKO mice (Fig 2D), further supporting no genotype differences in mitochondrial protein abundance. To determine whether there were functional impairments in mitochondrial respiration between genotypes, we performed high-resolution respirometry in permeabilized EDL muscle fibers. Based on a standard carbohydrate-driven SUIT protocol (Fig 2E), leak respiration, complex I (CI)-driven respiration, CI/complex II (CI+CII)-driven respiration and uncoupled respiration were not different between i-mPCKO and WT mice (Fig 2F).

**Figure 2.**
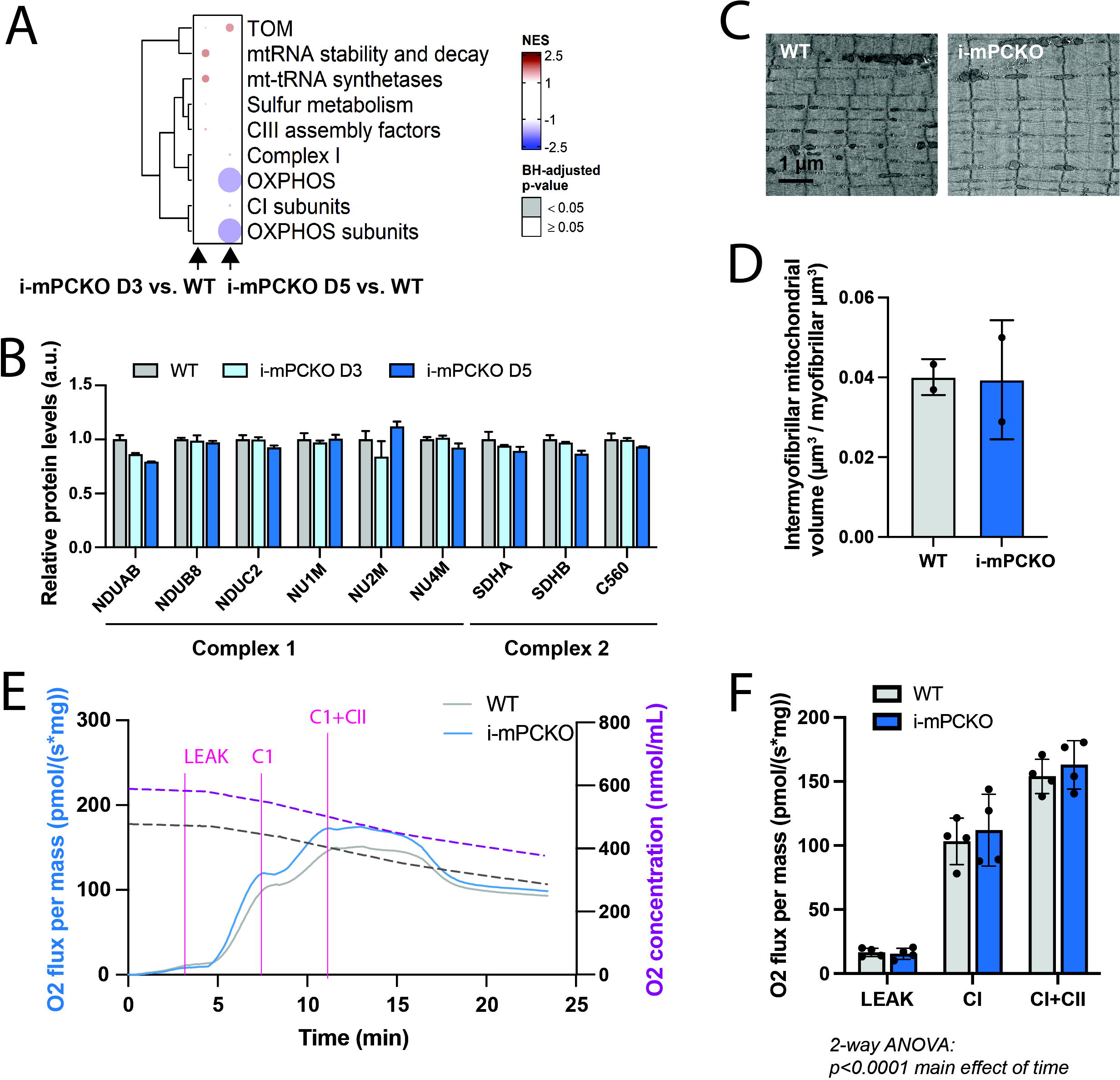
Mitochondrial protein abundance and respiratory function in normal in skeletal muscle from i-mPCKO mice. ***(A)*** Heatmap of FGSEA results comparing the abundance of proteins in MitoCarta pathways in skeletal muscle from male i-mPCKO (D3 and D5 post TMX) and WT mice (D5 post TMX) from publicly available proteomics data. Circles on the heatmap are colored by normalized enrichment score (NES) and scaled such that the lowest adjusted p-value is of maximum area. ***(B)*** Relative protein abundance of select proteins comprising complex I (CI) and complex II (CII) of the electron transport chain, from publicly available proteomics data in skeletal muscle from male i-mPCKO (D3 and D5 post TMX) and WT mice). ***(C)*** Representative (×5000 magnification) TEM images of intermyofibrillar mitochondria from male WT and i-mPCKO EDL muscle. ***(D)*** Quantification of the volume fraction of mitochondria/myofibrils (D5 post TMX). ***(E)*** Representative traces of oxygen flux overlayed with oxygen tension during cellular respirometry of permeabilized EDL fibers from WT and i-mPCKO mice (D5 post TMX; male). ***(F)*** Quantification of oxygen flux during leak, complex I (CI)-driven and complex I plus complex II-driven respiration (CI+CII) across all tested samples from WT and i-mPCKO mice (D5 post TMX; data are for male and female, which were collapsed together, for each genotype). Statistics: A, Gene-set enrichment analysis (GSEA). B, Student’s unpaired t-test. D, 2-way ANOVA with repeated measures across time, Šídák’s multiple comparisons test. Notation protein (*gene*): NDUAB (*Ndufa11*), NDUB8 (*Ndufb8*), NDUC2 (*Ndufc2*), NU1M (*Nd1*), NU2M (*Nd2*), NU4M (*Nd4*), SHDA (*Sdha*), SDHB (*Sdhb*), C560 (*Sdhc*).

### Permeabilization completely rescues contractile force in PCKO muscle fibers, indicating functional cross-bridge cycling

To determine whether actin-myosin and/or thin filament interactions were disrupted in i-mPCKO versus WT mice, we studied fibers from permeabilized soleus and EDL at D5 post initiation of TMX dosing. Permeabilization makes the sarcolemma porous, which enables experimental control over calcium and ATP concentrations via the bathing medium, and thus isolates cross-bridge cycling on regulated thin filaments from intracellular metabolic control and physiological calcium transients. Remarkably, when bathed in a high [Ca^2+^] solution with physiological levels of ATP, muscle fiber contractile function was completely rescued in i-mPCKO mice to the level of WT mice (Fig 3A-B). To determine whether thin filament sensitivity to calcium was different between genotypes, which would be masked at high [Ca^2+^], we bathed the fiber in solutions of varying [Ca^2+^] to generate a force-pCa curve. This rescue of force was not [Ca^2+^] dependent, as the force-pCa curves were indistinguishable between genotypes (Fig 3C), with calculated pCa_50_ values being similar between i-mPCKO and WT (Fig 3D). To further isolate whether there were changes in crossbridge cycling, we measured the speed of actomyosin contraction using an in vitro motility assay. Consistent with our results in permeabilized fibers, we observed no difference in speed between i-mPCKO and WT actomyosin (Fig 2E). Taken together, our results demonstrate that changes in i-mPCKO contraction are not due to alterations in actomyosin or thin filament proteins.

**Figure 3.**
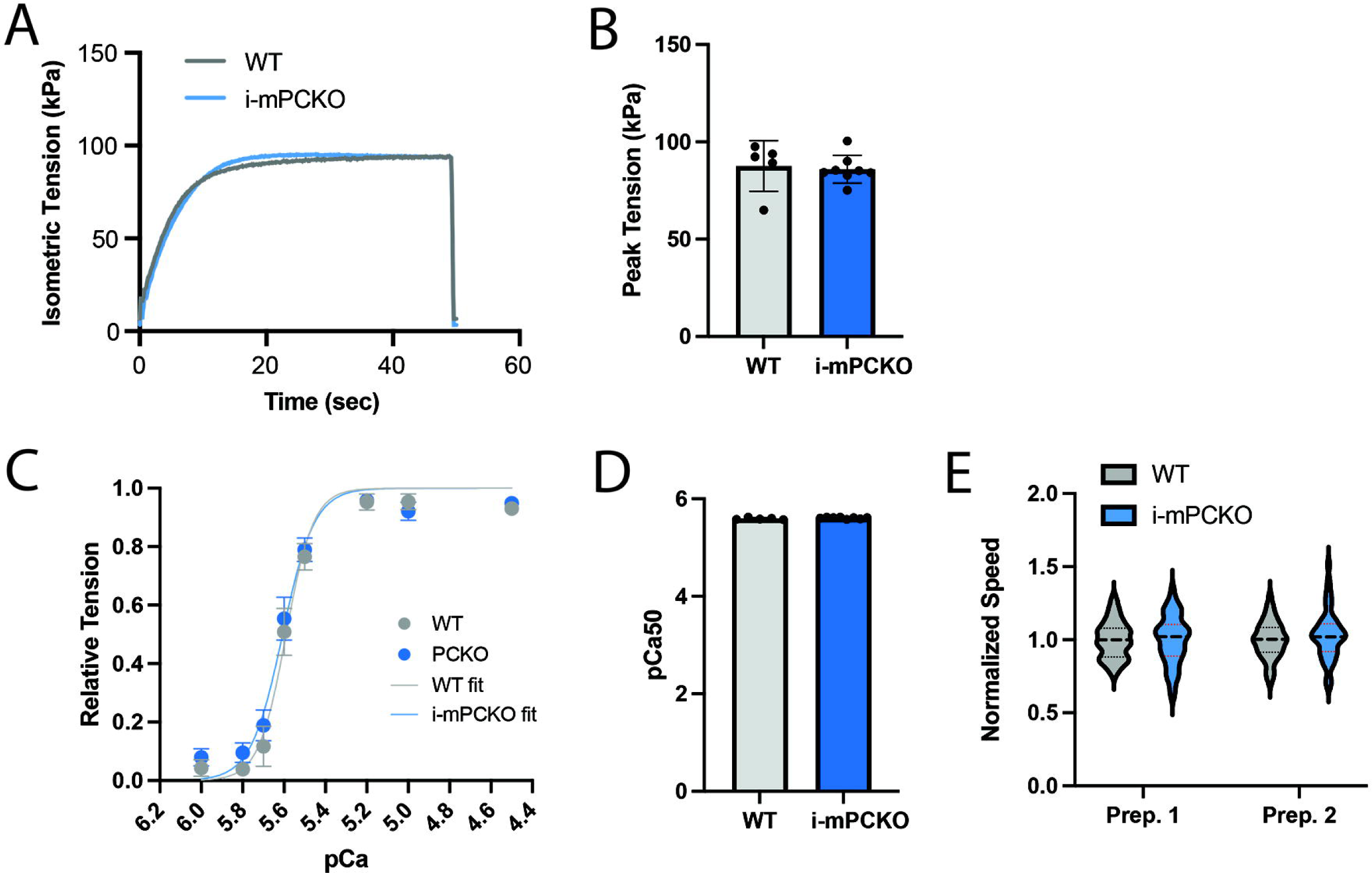
Normal cross-bridge cycling and myofilament motility in i-mPCKO mice. ***(A)*** Representative traces of isometric force elicited in permeabilized WT and i-mPCKO EDL fibers during immersion in a high Ca^2+^ (pCa 4.5) solution (D5 post TMX; male mice, only). ***(B)*** Quantification of peak tension of all fibers tested at pCa 4.5 (D5 post TMX; data are for male and female, which were collapsed together, for each genotype). ***(C)*** Force-pCa curves for WT and i-mPCKO permeabilized EDL fibers. Data for each EDL are fit with a Hill curve and the average fit is displayed for each genotype. (D5 post TMX; data are for male and female, which were collapsed together, for each genotype). ***(D)*** pCa50 values (pCa at which fitted curve reaches 0.5 of relative tension) for WT and i-mPCKO Hill cure fits. ***(E)*** Actin-myosin motility speed for two preparations (each preparation is pooled muscles from n=3 mice, with each datapoint representing the speed of an individual filament; 48-50 filaments were studied per genotype per prep). Statistics: B and D, Student’s unpaired t-test.

## Discussion

While reversible acetylation of lysine residues is a common and conserved post-translational modification that contributes to a diverse array of cellular processes, the contribution of lysine acetylation, and lysine acetyltransferases in particular, to skeletal muscle physiology and function are only beginning to be revealed. Building on previous work demonstrating that loss of p300 and CBP in skeletal muscle in adulthood dramatically impairs *ex vivo* contractile function in a matter of days, here we not only confirm this *ex vivo* phenotype, we demonstrate robust contractile dysfunction in i-mPCKO mice, *in vivo*. Mechanistically, the regulatory role of p300/CBP on skeletal muscle contractile function was not due to intrinsic issues in mitochondrial function, actomyosin cycling, or thin filament activation. In fact, contractile function in i-mPCKO mice was completely, and remarkably, recovered in permeabilized muscle. Taken together, these data implicate an important role of p300/CBP acetyltransferase activity in the regulation of skeletal muscle contractile function and suggest that regulation of contractile function by p300/CBP is via the mechanism(s) separate from the mitochondrial function and cross-bridge cycling.

Cross-bridge cycling through actin and myosin interaction forms the foundation of force development by skeletal muscle (25). Interestingly, many of the proteins involved in contractile function in skeletal muscle, including sarcomeric proteins supporting cross-bridge cycling and proteins involved in excitation-contraction (E-C) coupling are acetylated, thereby suggesting that acetylation may contribute to contractile function (2, 3, 26-28). A potential functional role for lysine acetylation in cross bridge cycling and skeletal muscle contractile function is supported by altered actin-myosin interactions with experimental manipulation of acetylation status of sarcomeric proteins across several model systems (9, 10, 29, 30). These include: 1) chemical acetylation of troponin C isolated from rabbit skeletal muscle increases Ca^2+^ sensitivity of skeletal myofibrils (9), 2) mimicking acetylation of actin in indirect-flight (skeletal) muscle in *Drosophila melanogaster* enhanced actomyosin associations (10), 3) acetylation of cardiac α- and β-myosin heavy chain (MHC) increased actin-sliding velocity (29), 4) increased pan protein acetylation increases cardiac myofilament Ca^2+^ sensitivity (31), 5) acetylation of muscle LIM protein and MHC positively correlates with cardiac contractile function (29), and 6) acetylation of lysine 132 on cardiac troponin I (cTnI) modulates myofilament relaxation and calcium sensitivity (30). While it is clear from these studies in skeletal and cardiac muscle that acetylation can regulate various aspects of actin-myosin interactions and calcium sensitivity, our findings demonstrate that contractile dysfunction in i-mPCKO mice is not through loss of integrity of cross-bridge cycling or thin filament calcium sensitivity since loss of contractile function in i-mPCKO muscle was *completely* rescued by simply permeabilizing the muscle fiber. Instead, our findings suggest that regulation of contractile function by p300 and CBP could be through effects on Ca^2+^ handling and/or ATP availability.

Through the actions of myosin ATPase, ATP is exclusively used by skeletal muscle to produce tension through cross-bridge cycling (25). Mitochondria are a primary source of ATP for contraction, especially during repeated contractions, making them fundamentally important to contractile function. In addition, because acetylomic analysis in skeletal muscle demonstrates that many mitochondrial and metabolic proteins are acetylated (2, 3), we investigated whether mitochondrial function was impaired in i-mPCKO mice. Interestingly, our data suggest that the contractile phenotype in i-mPCKO mice is also not due to deficits in the abundance of mitochondrial proteins (as determined from unbiased analysis of proteomic data), intermyofibrillar mitochondrial volume (as determined from TEM images), or impairments in mitochondrial function (as determined from high-resolution respirometry). Taken together, these findings assimilate with our recent data demonstrating that succinate dehydrogenase activity in the tibialis anterior (and diaphragm) is not different between i-mPCKO and WT mice (7), and infer that contractile dysfunction in i-mPCKO mice is not due to an inability of mitochondria to provide ATP.

A limitation of this work is that our assays in permeabilized fibers, be it for assessment of mitochondrial function or contractile function, are conducted in solutions in which we tightly control substrate and ionic concentrations; that is, our data demonstrate that mitochondrial respiratory function and cross-bridge function are functionally capable in an ‘ideal’ environment. It is possible, however, that the native environment in i-mPCKO muscle is dysregulated (e.g. low availability of metabolites or high concentrations of Mg^2+^), which in turn impacts mitochondrial function and/or cross-bridge cycling. While this is possible, we hypothesize that p300/CBP regulate contractile function through protein(s) that lie outside the mitochondria and sarcomere. For example, both SERCA1a and the ryanodine receptor (Ryr1) are acetylated on lysine residues in skeletal muscle (2), and p300-mediated lysine acetylation of lysine 514 of SERCA2a has been recently shown to regulate cardiomyocyte contractility and function (32), suggesting a mechanism by which acetylation could regulate Ca^2+^ release and reuptake. Furthermore, the cytoskeletal protein, tubulin, is acetylated on a residue that affects its association with membrane bound channels, suggesting a mechanism by which acetylation could regulate membrane excitability (33). A potential role for either of these mechanisms is supported by our permeabilized fiber data, in which contractile function was restored by providing Ca^2+^ and/or bypassing the membrane potential.

In conclusion, the importance of lysine acetylation to contractile function in skeletal muscle is only beginning to be fully elucidated. Here we establish the criticality of p300 and CBP to skeletal muscle contractile function, *in vivo*, and reaffirm our previous *ex vivo* work (7). Interestingly, our efforts to determine mechanisms of action reveal that neither mitochondrial abundance or respiration, or myosin and actin cross-bridge cycling are the underlying driver of the contractile phenotype in i-mPCKO mice. Considering the proteins involved in Ca^2+^ release and reuptake and membrane excitability are also acetylated, it will be interesting to determine in future work if p300/CBP regulate contractile function through one, or both, of these mechanisms.

## Acknowledgements

G.A.M. and S.S. were responsible for the conception and design of the study. G.A.M., S.S., J.L.A.F, A.G., J.A.S., K.S., L.S.F. and M.G. contributed to acquisition, analysis and interpretation of the data. G.A.M. and S.S. were responsible for the drafting of the manuscript and G.A.M., S.S., J.L.A.F, A.G., J.A.S., K.S., L.S.F. and M.G. all contributed to critical revision of the manuscript. All authors read and approved the final version of this manuscript and agreed to be accountable for all aspects of the work.

The authors thank Karen Shen for her assistance with troubleshooting the permeabilized fiber contractility assay.

## References

1. Choudhary C, Weinert BT, Nishida Y, Verdin E, and Mann M. The growing landscape of lysine acetylation links metabolism and cell signalling. Nat Rev Mol Cell Biol 15: 536–550, 2014.

2. Lundby A, Lage K, Weinert BT, Bekker-Jensen DB, Secher A, Skovgaard T, Kelstrup CD, Dmytriyev A, Choudhary C, Lundby C, and Olsen JV. Proteomic analysis of lysine acetylation sites in rat tissues reveals organ specificity and subcellular patterns. Cell Rep 2: 419–431, 2012.

3. Hostrup M, Lemminger AK, Stocks B, Gonzalez-Franquesa A, Larsen JK, Quesada JP, Thomassen M, Weinert BT, Bangsbo J, and Deshmukh AS. High-intensity interval training remodels the proteome and acetylome of human skeletal muscle. Elife 11: 2022.

4. Sebti S, Prébois C, Pérez-Gracia E, Bauvy C, Desmots F, Pirot N, Gongora C, Bach AS, Hubberstey AV, Palissot V, Berchem G, Codogno P, Linares LK, Liaudet-Coopman E, and Pattingre S. BAT3 modulates p300-dependent acetylation of p53 and autophagy-related protein 7 (ATG7) during autophagy. Proc Natl Acad Sci U S A 111: 4115–4120, 2014.

5. Shi D, Pop MS, Kulikov R, Love IM, Kung AL, and Grossman SR. CBP and p300 are cytoplasmic E4 polyubiquitin ligases for p53. Proc Natl Acad Sci U S A 106: 16275–16280, 2009.

6. Aslan JE, Rigg RA, Nowak MS, Loren CP, Baker-Groberg SM, Pang J, David LL, and McCarty OJ. Lysine acetyltransfer supports platelet function. J Thromb Haemost 13: 1908–1917, 2015.

7. Svensson K, LaBarge SA, Sathe A, Martins VF, Tahvilian S, Cunliffe JM, Sasik R, Mahata SK, Meyer GA, Philp A, David LL, Ward SR, McCurdy CE, Aslan JE, and Schenk S. p300 and cAMP response element-binding protein-binding protein in skeletal muscle homeostasis, contractile function, and survival. J Cachexia Sarcopenia Muscle 11: 464–477, 2020.

8. Martins VF, LaBarge SA, Stanley A, Svensson K, Hung CW, Keinan O, Ciaraldi TP, Banoian D, Park JE, Ha C, Hetrick B, Meyer GA, Philp A, David LL, Henry RR, Aslan JE, Saltiel AR, McCurdy CE, and Schenk S. p300 or CBP is required for insulin-stimulated glucose uptake in skeletal muscle and adipocytes. JCI Insight 7: 2022.

9. Grabarek Z, Mabuchi Y, and Gergely J. Properties of troponin C acetylated at lysine residues. Biochemistry 34: 11872–11881, 1995.

10. Viswanathan MC, Blice-Baum AC, Schmidt W, Foster DB, and Cammarato A. Pseudo-acetylation of K326 and K328 of actin disrupts Drosophila melanogaster indirect flight muscle structure and performance. Front Physiol 6: 116, 2015.

11. Schmidt W, Madan A, Foster DB, and Cammarato A. Lysine acetylation of F-actin decreases tropomyosin-based inhibition of actomyosin activity. J Biol Chem 295: 15527–15539, 2020.

12. McCarthy JJ, Srikuea R, Kirby TJ, Peterson CA, and Esser KA. Inducible Cre transgenic mouse strain for skeletal muscle-specific gene targeting. Skelet Muscle 2: 8, 2012.

13. Kang-Decker N, Tong C, Boussouar F, Baker DJ, Xu W, Leontovich AA, Taylor WR, Brindle PK, and van Deursen JM. Loss of CBP causes T cell lymphomagenesis in synergy with p27Kip1 insufficiency. Cancer Cell 5: 177–189, 2004.

14. Kasper LH, Fukuyama T, Biesen MA, Boussouar F, Tong C, de Pauw A, Murray PJ, van Deursen JM, and Brindle PK. Conditional knockout mice reveal distinct functions for the global transcriptional coactivators CBP and p300 in T-cell development. Mol Cell Biol 26: 789–809, 2006.

15. Chleboun GS, Patel TJ, and Lieber RL. Skeletal muscle architecture and fiber-type distribution with the multiple bellies of the mouse extensor digitorum longus muscle. Acta Anat (Basel) 159: 147–155, 1997.

16. Roche SM, Gumucio JP, Brooks SV, Mendias CL, and Claflin DR. Measurement of Maximum Isometric Force Generated by Permeabilized Skeletal Muscle Fibers. J Vis Exp e52695, 2015.

17. Kalakoutis M, Di Giulio I, Douiri A, Ochala J, Harridge SDR, and Woledge RC. Methodological considerations in measuring specific force in human single skinned muscle fibres. Acta Physiol (Oxf) 233: e13719, 2021.

18. Walker JS, Li X, and Buttrick PM. Analysing force-pCa curves. J Muscle Res Cell Motil 31: 59–69, 2010.

19. Rath S, Sharma R, Gupta R, Ast T, Chan C, Durham TJ, Goodman RP, Grabarek Z, Haas ME, Hung WHW, Joshi PR, Jourdain AA, Kim SH, Kotrys AV, Lam SS, McCoy JG, Meisel JD, Miranda M, Panda A, Patgiri A, Rogers R, Sadre S, Shah H, Skinner OS, To TL, Walker MA, Wang H, Ward PS, Wengrod J, Yuan CC, Calvo SE, and Mootha VK. MitoCarta3.0: an updated mitochondrial proteome now with sub-organelle localization and pathway annotations. Nucleic Acids Res 49: D1541–D1547, 2021.

20. West MJ. Estimating volume in biological structures. Cold Spring Harb Protoc 2012: 1129–1139, 2012.

21. KIELLEY WW, and BRADLEY LB. The relationship between sulfhydryl groups and the activation of myosin adenosinetriphosphatase. J Biol Chem 218: 653–659, 1956.

22. Spudich JA, and Watt S. The regulation of rabbit skeletal muscle contraction. I. Biochemical studies of the interaction of the tropomyosin-troponin complex with actin and the proteolytic fragments of myosin. J Biol Chem 246: 4866–4871, 1971.

23. Clippinger SR, Cloonan PE, Greenberg L, Ernst M, Stump WT, and Greenberg MJ. Disrupted mechanobiology links the molecular and cellular phenotypes in familial dilated cardiomyopathy. Proc Natl Acad Sci U S A 116: 17831–17840, 2019.

24. Barrick SK, Greenberg L, and Greenberg MJ. A troponin T variant linked with pediatric dilated cardiomyopathy reduces the coupling of thin filament activation to myosin and calcium binding. Mol Biol Cell 32: 1677–1689, 2021.

25. Huxley AF. Cross-bridge action: present views, prospects, and unknowns. J Biomech 33: 1189–1195, 2000.

26. Liang D, Chen C, Huang S, Liu S, Fu L, and Niu Y. Alterations of Lysine Acetylation Profile in Murine Skeletal Muscles Upon Exercise. Front Aging Neurosci 14: 859313, 2022.

27. Huang S, Liu S, Niu Y, and Fu L. Scriptaid/exercise-induced lysine acetylation is another type of posttranslational modification occurring in titin. J Appl Physiol (1985) 128: 276–285, 2020.

28. Ryder DJ, Judge SM, Beharry AW, Farnsworth CL, Silva JC, and Judge AR. Identification of the Acetylation and Ubiquitin-Modified Proteome during the Progression of Skeletal Muscle Atrophy. PLoS One 10: e0136247, 2015.

29. Samant SA, Pillai VB, Sundaresan NR, Shroff SG, and Gupta MP. Histone Deacetylase 3 (HDAC3)-dependent Reversible Lysine Acetylation of Cardiac Myosin Heavy Chain Isoforms Modulates Their Enzymatic and Motor Activity. J Biol Chem 290: 15559–15569, 2015.

30. Lin YH, Schmidt W, Fritz KS, Jeong MY, Cammarato A, Foster DB, Biesiadecki BJ, McKinsey TA, and Woulfe KC. Site-specific acetyl-mimetic modification of cardiac troponin I modulates myofilament relaxation and calcium sensitivity. J Mol Cell Cardiol 139: 135–147, 2020.

31. Eaton DM, Martin TG, Kasa M, Djalinac N, Ljubojevic-Holzer S, Von Lewinski D, Pöttler M, Kampaengsri T, Krumphuber A, Scharer K, Maechler H, Zirlik A, McKinsey TA, Kirk JA, Houser SR, Rainer PP, and Wallner M. HDAC Inhibition Regulates Cardiac Function by Increasing Myofilament Calcium Sensitivity and Decreasing Diastolic Tension. Pharmaceutics 14: 2022.

32. Gorski PA, Lee A, Lee P, Oh JG, Vangheluwe P, Ishikawa K, Hajjar R, and Kho C. Identification and Characterization of p300-Mediated Lysine Residues in Cardiac SERCA2a. Int J Mol Sci 24: 2023.

33. Zampar GG, Chesta ME, Carbajal A, Chanaday NL, Díaz NM, Casale CH, and Arce CA. Acetylated tubulin associates with the fifth cytoplasmic domain of Na(+)/K(+)-ATPase: possible anchorage site of microtubules to the plasma membrane. Biochem J 422: 129–137, 2009.

